# Discovery and functional annotation of quantitative trait loci affecting resistance to sea lice in Atlantic salmon

**DOI:** 10.1101/455626

**Authors:** Diego Robledo, Alejandro P. Gutiérrez, Agustín Barría, Jean P. Lhorente, Ross D. Houston, José M. Yáñez

## Abstract

Sea lice (*Caligus rogercresseyi*) are ectoparasitic copepods which have a large negative economic and welfare impact in Atlantic salmon (*Salmo salar*) aquaculture, particularly in Chile. A multi-faceted prevention and control strategy is required to tackle lice, and selective breeding contributes via cumulative improvement of host resistance to the parasite. While host resistance has been shown to be heritable, little is yet known about the individual loci that contribute to this resistance, the potential underlying genes, and their mechanisms of action. In this study we took a multifaceted approach to identify and characterise quantitative trait loci (QTL) affecting hose resistance in a population of 2,688 Caligus-challenged Atlantic salmon post-smolts from a commercial breeding programme. We used low and medium density genotyping to collect genome-wide SNP marker data for all animals. Moderate heritablility estimates of 0.28 and 0.24 were obtained for lice density (as a measure of host resistance) and growth during infestation respectively. Three QTL explaining between 7 and 13 % of the genetic variation in resistance to sea lice (as represented by the traits of lice density) were detected on chromosomes 3, 18 and 21. Characterisation of these QTL regions was undertaken using RNA sequencing and pooled whole genome sequencing data. This resulted in the identification of a shortlist of potential underlying causative genes, and candidate functional mutations for further study. For example, candidates within the chromosome 3 QTL include a putative premature stop mutation in TOB1 (an anti-proliferative transcription factor involved in T cell regulation) and an uncharacterized protein which showed significant differential allelic expression (implying the existence of a cis-acting regulatory mutation). While host resistance to sea lice is polygenic in nature, the results of this study highlight significant QTL regions together explaining a moderate proportion of the heritability of the trait. Future investigation of these QTL may enable improved knowledge of the functional mechanisms of host resistance to sea lice, and incorporation of functional variants to improve genomic selection accuracy.

## 1. INTRODUCTION

Sea lice are a major concern for salmon aquaculture worldwide, in particular *Lepeophtheirus salmonis* in the Northern Hemisphere and *Caligus rogercresseyi* in the Southern, and cause losses of over $430M per year (Costello 2009). These copepods attach to the skin and feed on the mucus and blood of several species of salmonid fish. Parasitized fish display reduced growth rate and increased occurrence of secondary infections (Jónsdóttir et al. 1992). In addition to a significant negative impact on salmonid health and welfare, lice prevention and treatment costs are a large economic burden for salmonid aquaculture. Current control strategies include, for example, feed supplements, cleaner fish, tailored cage design, or ‘lice-zapping’ lasers (Aaen et al. 2015), but these multifaceted strategies are only partially effective. Expensive and potentially environmentally damaging chemicals and treatments are still frequently required to control sea lice populations, which are becoming resistant to common delousing drugs (Bravo et al. 2008; Aaen et al. 2015). Therefore, despite these extensive control efforts, sea lice remain a significant threat to salmon welfare and aquaculture sustainability, and incur in further indirect costs via negative impact on public opinion of aquaculture (Jackson et al. 2017).

Selective breeding can contribute to sea lice prevention via harnessing naturally occurring genetic variation within commercial salmon stocks to identify the most resistant individuals. The identification of selection candidates can be enabled either by pedigree or genomic based approaches, the latter via genomic selection (Meuwissen et al. 2001). Moderate genetic variation in resistance to sea lice exists in Atlantic salmon, with heritabilities ranging between 0.1 and 0.3 for both the North Atlantic sea louse (*Lepeophtheirus salmonis*; Kolstad et al. 2005; Gjerde et al. 2011; Ødegård et al. 2014; Gharbi et al. 2015; Tsai et al. 2016), and the Pacific sea louse (*Caligus rogercresseyi*; Lhorente et al. 2012; Yáñez et al. 2014; Correa et al. 2017a); and genomic selection approaches for resistance against both lice species have yielded substantially higher prediction accuracies that ‘traditional’ pedigree-based approaches (Ødegård et al. 2014; Gharbi et al. 2015; Tsai et al. 2016; Correa et al. 2017b).

Genomic selection is now routinely applied in Atlantic salmon breeding programmes for the genetic improvement of several traits (Houston 2017; Lhorente et al. Submitted). While it offers notable benefits in terms of selection accuracy, it has limitations such as the significant cost (via the need for high volume genotyping using SNP arrays), and the limited accuracy when the reference and selection candidate populations are not closely related (Daetwyler et al. 2012; Tsai et al. 2016). Discovering the causative genetic polymorphisms underlying phenotypic variation in complex traits is a fundamental goal of genetic research. Further, identifying these causative variants would also facilitate more effective genomic selection, potentially via cheaper genoyping strategies, increased genetic gain each generation, and improved persistency of prediction accuracy across generations and populations. Further, knowledge of causative variants offer the future possibilities of harnessing genomic editing approaches, for example to introduce / remove favourable / detrimental genetic variants. However, finding causative mutations within QTL regions is very challenging, with few success stories in farm animals to date. QTL regions tend to cover large segments of chromosomes, and typically contain many variants in linkage disequilibrium that show approximately equal association with the trait. Functional genome annotation data can be applied to prioritise variants within these regions and – although largely lacking for aquaculture species to date – are currently being developed for Atlantic salmon and other salmonid species (Macqueen et al. 2017). While laborious, shortlisting and identification of causative genes and variants impacting on disease resistance has positive implications for both selective breeding, and fundamental understanding of the host-pathogen interaction.

In the current study, a large population of farmed Atlantic salmon of Chilean origin was challenged with sea lice and high density SNP genotype data was collected. The overall aim of the study was to detect and annotate QTL affecting host resistance to sea lice in farmed Atlantic salmon, with a view to identifying putative causative genes and variants. The specific objectives were a) estimate genetic variance of sea lice resistance in our population, b) dissect the genetic architecture of the trait, and c) explore the genomic basis of the detected QTL using transcriptomics and whole-genome sequencing data.

## 2. METHODS

### 2.1. Disease challenge

2,668 Atlantic salmon (*Salmo salar*) Passive Integrated Transponder (PIT)-tagged post-smolts (average weight 122 g, measured for weight and length prior to the start of the challenge) from 104 families from the breeding population of AquaInnovo (Salmones Chaicas, Xth Region, Chile), were experimentally challenged with *Caligus rogercresseyi* (chalimus II-III). Infestation with the parasite was carried out by using 50 copepods per fish and stopping the water flow for 6 h during infestation, thereafter water flow was gradually restored reaching its normal flow 2 days after. During this process oxygen saturation was maintained at 90-110 %, and oxygen and temperature where constantly monitored. Eight days after the infestation fish were sedated, carefully and individually removed from the tanks, and the number of sea lice attached where counted from head to tail. At this stage, fish were measured for weight and length, pit-tags were read, and fin-clips collected for DNA extraction. Log-transformed lice density was estimated as *log_e_* (*LC* / *BW*_ini_ ^2/3^), where LC is the number of lice counted on the fish, BW_ini_ is the initial body weight prior to the challenge, and BW_ini_^2/3^ is an approximation of the surface of the skin of each fish (Ødegård et al. 2014). Growth during infestation was calculated as [(BW_end_ - BW_ini_) / BW_ini_) * 100], where BW_ini_ and BW_end_ are the weight of the fish at the start and at the end of the trial respectively; the same formulae was used for length.

### 2.2 Genotyping

DNA was extracted from fin-clips from challenged fish using a commercial kit (Wizard^®^ Genomic DNA Purification Kit, Promega), following the manufacturer’s instructions. All samples where genotyped with a panel of 968 SNPs chosen as a subset of the SNPs from a medium density SNP array (Yáñez et al. 2016; Supplementary Table 1) using Kompetitive Allele Specific PCR (KASP) assays (LGC Ltd, UK). A population containing full-siblings of the challenged animals had previously been genotyped with a SNP panel of 45,819 SNPs (n = 1,056, Correa et al. 2015; Yáñez et al. 2016), and the experimental lice-challenged population was imputed to ~46 K SNPs using FImpute v.2.2 (Sargolzaei et al. 2014). Imputation accuracy was estimated by 10-fold cross validation, masking 10% of the 1,056 genotyped full-sibs to the 968 SNP panel, performing imputation, and then assessing the correlation between the true genotypes and the imputed genotypes. All imputed SNPs showing imputation accuracy below 80% were discarded. Genotypes were further filtered and removed according to the following criteria: SNP call-rate < 0.9, individual call-rate < 0.9, FDR rate for high individual heterozygosity < 0.05, identity-by-state > 0.95 (both individuals removed), Hardy-Weinberg equilibrium p-value < 10^–6^, minor allele frequency < 0.01. After filtering 38,028 markers and 2,345 fish remained for association analysis.

### 2.3 Estimation of genetic parameters

Variance components and heritabilities were estimated by ASReml 4.1 (Gilmour *et al.* 2014) fitting the following linear mixed model:

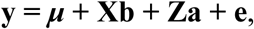

where **y** is a vector of observed phenotypes (lice number, lice density, initial weight, initial length and weight and length gain during infestation), *μ* is the overall mean of phenotype records, **b** is the vector of fixed effects of tank (as factor with 3 levels) and initial body weight (as a covariate; except when initial weight or initial length were the observed phenotypes), **a** is a vector of random additive genetic effects of the animal, distributed as ~N(0,**G***σ*^2^a) where *σ*^2^a is the additive (genetic) variance, **G** is the genomic relationship matrix. **X** and **Z** are the corresponding incidence matrices for fixed and additive effects, respectively, and **e** is a vector of residuals. The best model was determined comparing the log-likehood of models with different fixed effects and covariates. Phenotypic sex was not significant for any of the traits. **G** was calculated using the GenABEL R package (Aulchenko *et al.* 2007) to obtain the kinship matrix using the method of Amin et al. (2007), which was multiplied by a factor of 2 and inverted using a standard R function. Genetic correlations were estimated using bivariate analyses implemented in ASReml 4.1 (Gilmour *et al.* 2014) fitting the same fixed and random effects described in the univariate linear mixed model described above.

### 2.4 Single-SNP genome-wide association study

The single-SNP GWAS was performed using the GenABEL R package (Aulchenko *et al.* 2007) by applying the mmscore function (Chen and Abecasis, 2007), which accounts for the relatedness between individuals applied through the genomic kinship matrix. Significance thresholds were calculated using a Bonferroni correction where genome-wide significance was defined as 0.05 divided by number of SNPs (Duggal *et al.* 2008) and suggestive as one false positive per genome scan (1 / number SNPs).

### 2.5 Regional heritability mapping

A regional heritability mapping (RHM) analysis (Nagamine *et al.* 2012; Uemoto *et al.* 2013) was performed where the genome was divided into overlapping regions consisting of 50 sequential SNPs and overlapping by 25 SNPs using Dissect v.1.12.0 (Canela-Xandri *et al.* 2015). The significance of the regional heritability for each window was evaluated using a log likelihood ratio test statistic (LRT) comparing the global model fitting all markers with the model only fitting SNPs in a specific genomic region.

### 2.6 Whole-genome sequencing

Genomic DNA from a pool of 50 fish with high sea lice counts (Mean = 48) and a pool of 50 fish with low sea lice counts (Mean = 20) were sequenced in five lanes of HiSeq 4000 as 150 bp PE reads. Family structure was similar in both pools, with 34 different families and a maximum of two fish per family. The quality of the sequencing output was assessed using FastQC v.0.11.5 (http://www.bioinformatics.babraham.ac.uk/projects/fastqc/). Quality filtering and removal of residual adaptor sequences was conducted on read pairs using Trimmomatic v.0.32 (Bolger et al. 2004). Specifically, Illumina adaptors were clipped from the reads, leading and trailing bases with a Phred score less than 20 were removed, and the read trimmed if the sliding window average Phred score over four bases was less than 20. Only reads where both pairs were longer than 36 bp post-filtering were retained. Filtered reads were mapped to the most recent Atlantic salmon genome assembly (ICSASG_v2; Genbank accession GCF_000233375.1; Lien et al. 2016) using Burrows-Wheeler aligner v.0.7.8 BWA-MEM algorithm (Li 2013). Pileup files describing the base-pair information at each genomic position were generated from the alignment files using the mpileup function of samtools v1.4 (Li et al. 2009), discarding those aligned reads with a mapping quality < 30 and those bases with a Phred score < 30. Synchronized files containing read counts for every allele variant in every position of the genome were obtained using Popoolation2 v1.201 (Kofler et al. 2011). A read depth ≥ 10 and a minimum of 3 reads of the minor allele were required for SNP calling.

### 2.7 Differential allelic expression

The sequence data from a previous RNA-Seq study on the skin of animals of this sea lice infected population (Robledo et al. 2018) were used to investigate allelic specific expression. Alignment files were produced using STAR v.2.5.2b (Dobin et al. 2013; detailed protocol can be found in Robledo et al. 2018) and used for SNP identification and genotype calling with samtools v1.4 (Li et al. 2009). Reads with mapping quality < 30 and bases with phred quality scores < 30 were excluded. A read depth ≥ 10 and ≥ 3 reads for the alternative allele were required to call a SNP. Read counts were obtained for each allele in heterozygous loci and a binomial test was performed to assess the significance of the allelic differences using the R package AllelicImbalance (Gådin et al. 2015).

## 3. RESULTS

### 3.1 Disease challenge, genotyping and imputation

Eight days after the start of the challenge, the average lice burden per fish across the challenged population was 38 ± 16. The average weight prior to the start of the trial was 122 ± 40 grams and 143 ± 49 grams after the challenge. All samples were genotyped using a low-density SNP panel (968 SNPs), but 50 samples were genotyped for less than 90 % of the SNPs and therefore removed from subsequent analyses. The remaining samples were imputed to high-density from a population of 1,056 salmon that included full siblings of the challenge population, which had previously been genotyped for 45K SNPs (subset of Yáñez et al. 2016 selected as described in Correa et al. 2015). After removing SNPs showing low imputation accuracy (< 80%), a total of 39,416 SNPs remained with an average imputation accuracy (as assessed by cross-validation) of ~95%. After further call rate, minimum allele frequency, heterozygosity, identity-by-descent and Hardy-Weinberg filters, 38,028 markers and 2,345 fish remained for downstream analyses.

### 3.2 Genetic parameters

Heritabilities and genetic correlations of different traits related to sea lice load, growth and growth during infestation are shown in Table 1. The estimated heritability for sea lice load was 0.29 ± 0.04, and the number of sea lice attached to each fish showed positive genetic correlation with both initial weight (0.47 ± 0.07) and initial length (0.42 ± 0.08). However, sea lice density (h^2^ = 0.28 ± 0.04) was independent of the size of the fish which implies that these positive genetic correlations are due to the fact that larger fish tend to have more lice. Initial weight and length showed significant heritabilities as expected, and growth during infestation also presented a moderate heritability, especially weight (h^2^ = 0.24 ± 0.04). Surprisingly, weight gain during infestation showed a positive genetic correlation with sea lice density and sea lice counts, albeit with a high standard error (0.25 ± 0.12 and 0.27 ± 0.12 respectively).

**Table 1.** Heritabilities and genetic correlations

|  | Caligus<br>number | Caligus<br>density<br>(Ln) | Initial<br>weight | Weight<br>gain (%) | Initial<br>length | Length<br>gain (%) |
| --- | --- | --- | --- | --- | --- | --- |
| **Caligus**<br>**number** | **$0.29 \pm 0.04$** |  |  |  |  |  |
| **Caligus**<br>**density (Ln)** | $0.78 \pm 0.04$ | **$0.28 \pm 0.04$** | | | | |
| **Initial weight** | $0.47 \pm 0.07$ | $-0.15 \pm 0.09$ | **$0.41 \pm 0.04$** | | | |
| **Weight gain**<br>**(%)** | $0.27 \pm 0.12$ | $0.25 \pm 0.12$ | $0.05 \pm 0.10$ | **$0.24 \pm 0.04$** | | |
| **Initial length** | $0.42 \pm 0.08$ | $-0.22 \pm 0.10$ | $0.99 \pm 0.01$ | $0.17 \pm 0.11$ | **$0.27 \pm 0.03$** | |
| **Length gain**<br>**(%)** | $0.23 \pm 0.15$ | $0.31 \pm 0.15$ | $0.09 \pm 0.03$ | $0.89 \pm 0.05$ | $0.09 \pm 0.15$ | **$0.12 \pm 0.03$** |

### 3.3 Genome-wide association

The genetic architectures for the traits of sea lice density (Figure 1) and growth gain during infestation (Supplementary Figure 1) were studied using two different methods. The single SNP GWAS for sea lice density revealed three SNPs reaching genome-wide significance in the distal part of chromosome 3 (Figure 1A), each estimated to explain 3.61 – 4.14 % of the genetic variation. Additional SNPs showed suggestive association with sea lice density in chromosome 5, chromosome 9 and chromosome 18. Regional heritability analyses using 50 SNP windows confirmed the QTL in chromosomes 3 and 18, both estimated to explain ~7.5 % of the genetic variation in sea lice density (Figure 1B). The RHM approach detected an additional QTL not found in the single SNP GWAS, on chromosome 21, explaining close to 10 % of the genetic variation in sea lice density. The QTL regions in chromosomes 3, 18 and 21 were further refined, adding and removing SNPs until the window explaining the most genetic variation for each QTL was found. These QTL were narrowed to regions of 3-5 Mb and were each estimated to explain between 7.8 and 13.4 % of the genetic variation in sea lice density, accounting in total for almost 30 % of the genetic variance (Table 2) assuming additive effects of the QTL.

**Figure 1.**
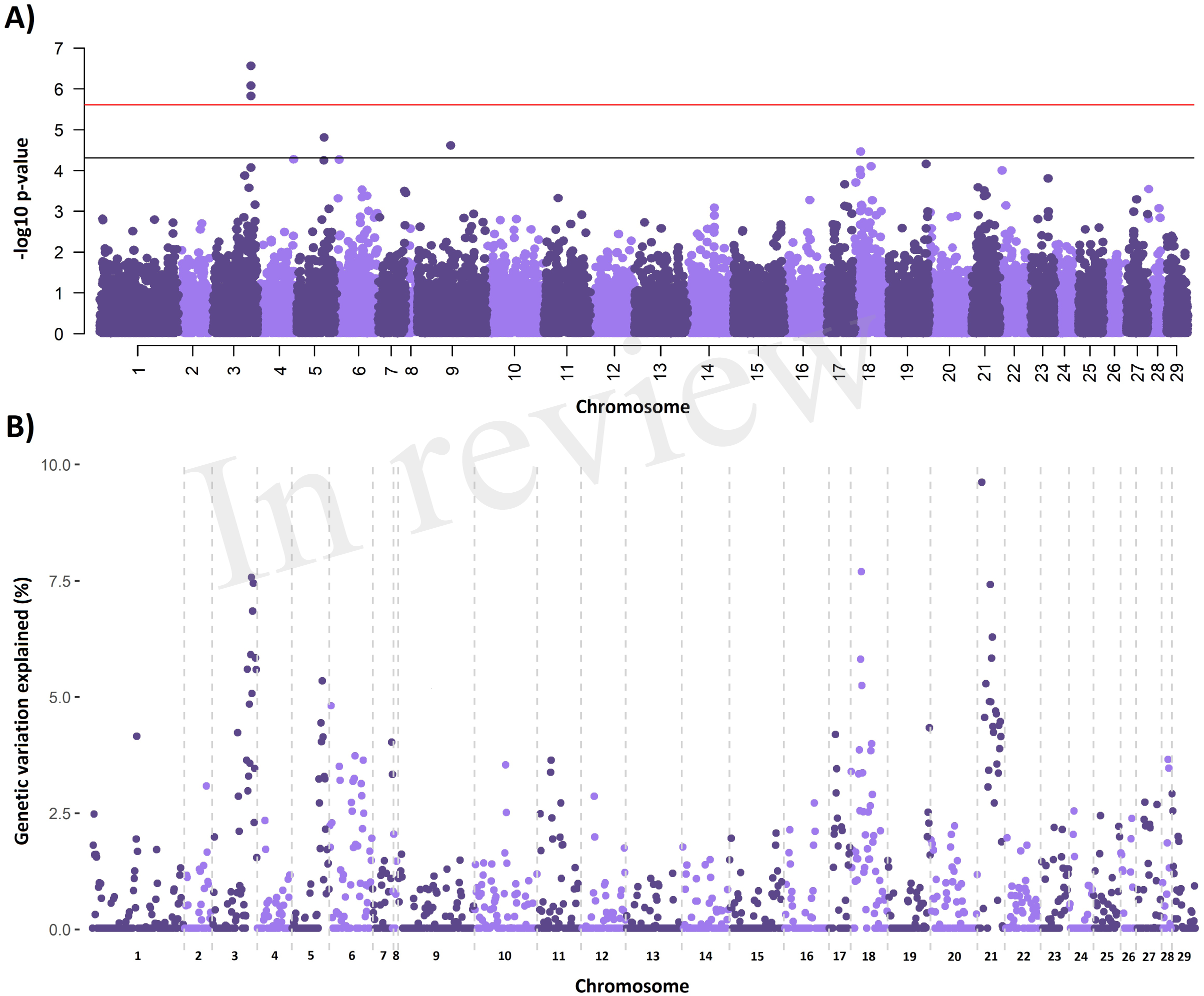
Genome-wide association analyses for sea lice density. A) Single-SNP GWAS results, horizontal bars represent Bonferroni corrected significance (red) and nominal significance (black). B) RHM results showing the percentage of additive genetic variance explained by each genomic region, represented by 50 consecutive SNPs.

**Table 2.** Details of Sea lice resistance QTL

| Chromosome | Start (Mb) | End (Mb) | Number of SNPs | Genetic variance explained (%) |
| --- | --- | --- | --- | --- |
| 3 | 77.54 | 82.54 | 58 | 7.82 |
| 18 | 14.87 | 17.56 | 62 | 8.34 |
| 21 | 7.96 | 10.44 | 52 | 13.39 |

Positions and details of the QTL detected by RHM. Chromosome, start and end boundaries of the QTL region, number of SNPs in the QTL region, and percentage of genetic variance of sea lice density explained by the QTL are shown.

### 3.4 QTL characterization

The three sea lice density QTL regions were then further interrogated to identify and characterise potential causative genes and variants. The Atlantic salmon genome annotation (Lien et al. 2016), the results of a previous RNA-Seq study comparing lice-attachment sites and healthy skin (Robledo et al. 2018), and SNP variants obtained from the WGS of pools of fish with high and low number of lice were combined to obtain a holistic view of these regions (Figure 2, Supplementary Figures 1 and 2). The QTL regions were all relatively large, and contained a large number of SNPs and genes. We detected 16 K – 22 K putative SNPs in each of the QTL regions, but less than a thousand were located in genic regions in each of them. Surprisingly, the number mutations that were predicted to have a moderate or large functional effect was relatively high, especially in chromosome 3 where a total of 213 non-synonymous SNPs were detected, along with 5 premature stop, 1 stop lost and 12 start gain mutations. The equivalent numbers were more modest for chromosomes 18 and 21, with 37 and 13 non-synonymous mutations respectively, but still relatively high to single out high priority candidate causative variants using variant effect prediction data alone. The three QTL regions had also a relatively high number of genes (83, 36 and 11 genes in the QTL regions of chromosomes 3, 18 and 21 respectively). Therefore, to shortlist candidate genes and variants, a combination of differential expression between resistant and susceptible fish, variant effect prediction, and a literature search relating to the function of the genes and their potential role in host response to parasitic infection were used.

**Figure 2.**
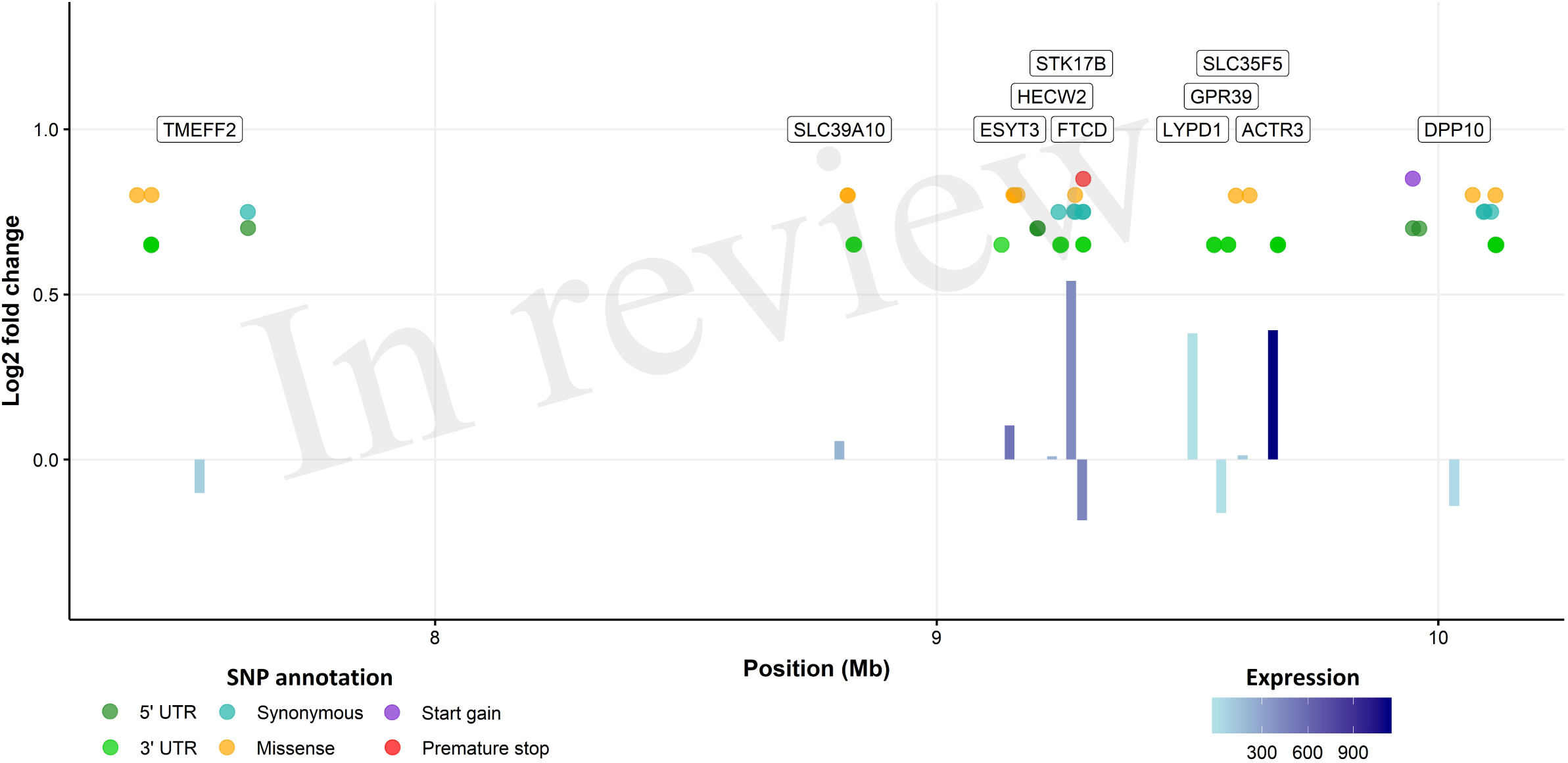
QTL region in chromosome 21. Bars represent the log2 fold change between healthy and sea lice attachment skin for every gene in the QTL region according to the RNA-seq. Bar colour represents the expression level of the gene (lighter = less expressed), and the annotation of the gene is presented in a label on the top of the graph. Genic SNPs detected by WGS are shown in between, those with putatively more severe effects are located towards the top of the figure.

The clearest candidate gene in chromosome 3 is TOB1, where a premature stop mutation was detected. This transcription factor negatively regulates cell proliferation (Matsuda et al. 1996), including that of T-cells (Baranzini 2014). In our study, it was highly expressed in the skin according to RNA-Seq data, and its expression was significantly lower in lice attachment regions of the skin (Robledo et al. 2018). For chromosome 21, serine / threnonine-protein kinase 17B (STK17B) showed the highest fold change between lice-attachment and healthy skin and a missense mutation; this gene has been connected to apoptosis and T-cell regulation, and T-cells of STK17B deficient mice are hypersensitive to stimulation (Honey 2005). Previous studies comparing the immune response of resistant and susceptible salmonid species have linked Th2-type responses to sea lice resistance (Braden et al. 2015), which is consistent with these two genes potentially having a functional role relating to the resistance QTL. Chromosome 18 does not contain any clear candidate genes, but from a literature search alone, the most plausible gene is probably heme binding protein 2 (HEBP2). Reducing iron availability has been suggested as a possible mechanism of resistance to sea lice (Fast et al. 2007; Sutherland et al. 2014) and *Piscirickettsia salmonis* (Pulgar et al. 2015) in Atlantic salmon.

Since complex traits can be influenced by causative mutations in regulatory regions that impact gene expression (Keane et al. 2011; Albert and Kruglyak, 2015) a differential allelic expression analysis was performed to screen for potential cis-acting regulatory mutations affecting genes in the QTL regions. An uncharacterized gene (XP_014049605.1) showed clear signs of differential allelic expression (P = 0.00081, Figure 3) in chromosome 3 at 8.1 Mb, less than 200 Kb away from the significant GWAS SNPs. This gene is also highly expressed in the skin of lice-infected salmon (Robledo et al. 2018), and similar proteins are found in other salmonid and teleost species.

**Figure 3.**
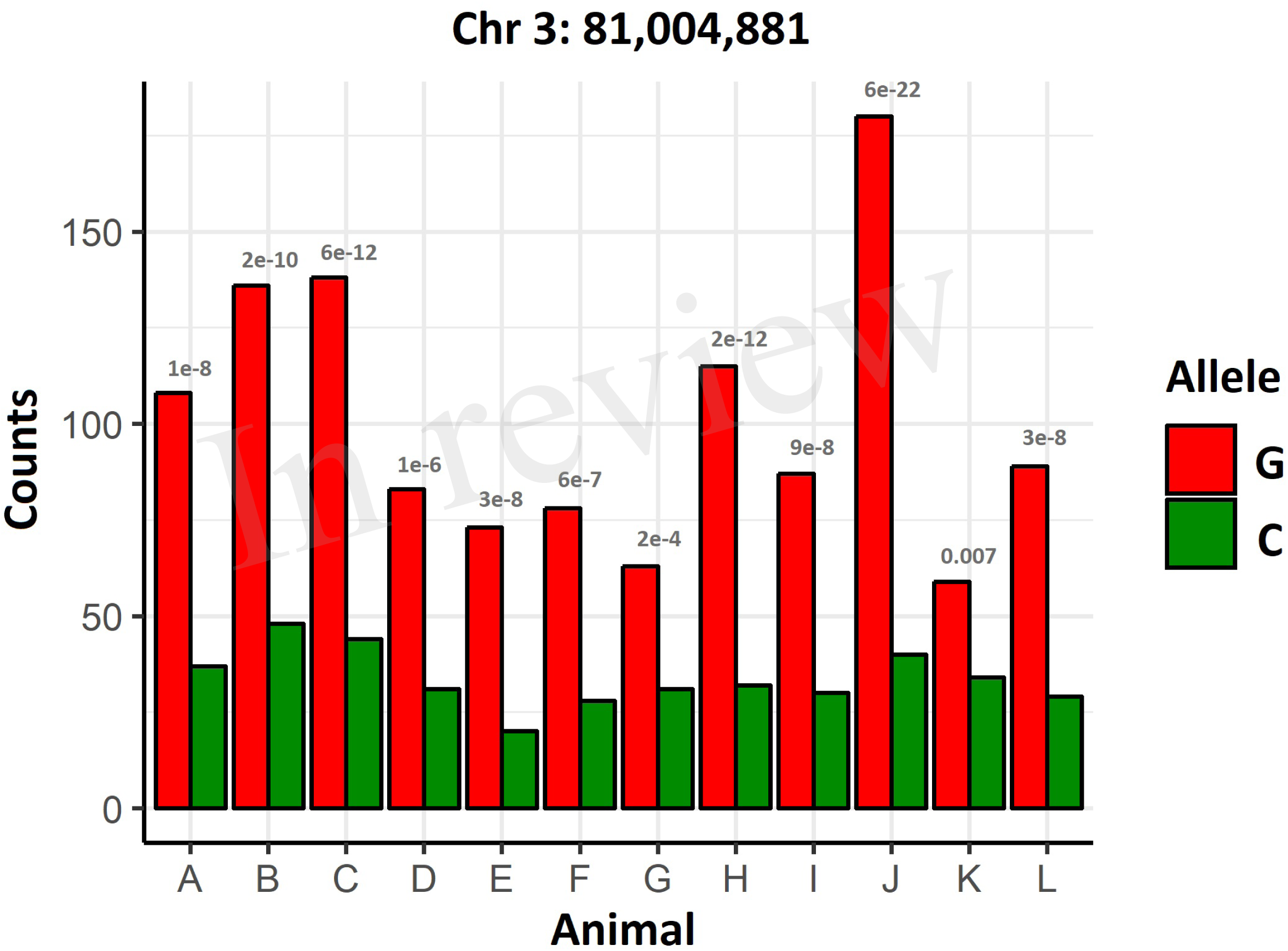
Differential allelic expression of XP_014049605.1. Gene counts in heterozygous animals for a SNP showing allelic imbalance in XP 014049605.1. The p-values of the binomial test comparing the expression of both alleles are shown for each animal.

## 4. DISCUSSION

In the current study we aimed to use a combination of GWAS, RNA-Seq, whole genome resequencing, and functional annotation approaches to identify and characterise QTL influencing host resistance to sea lice. Heritability estimates for sea lice density were similar or higher than previous studies on *C. rogercresseyi*-challenged Atlantic salmon. Lhorente et al. (2012, 2014) obtained pedigree-estimated heritabilities of 0.17 – 0.34, while estimates on a previously related sea lice challenged population were of 0.10 – 0.11 with both pedigree and molecular information (Yáñez et al., 2014; Correa et al., 2017a,b). This heritability is also in consistent with heritability estimates for salmon challenged with *L. salmonis* (0.2 – 0.3; Kolstad et al. 2005; Gjerde et al. 2011; Gharbi et al. 2015; Tsai et al. 2016), and similar to heritabilities for resistance to other ectoparasites affecting Atlantic salmon such as *Gyrodactylus salaris* (0.32; Salte et al. 2010) and *Neoparamoeba perurans* (Amoebic gill disease; 0.23 – 0.48; Taylor et al. 2009; Robledo et al. 2018).

Previous studies on the architecture of resistance to *C. rogercresseyi* had revealed just one significant SNP in chromosome 21 ~6.5 Mb (Correa et al. 2017a). While no significant SNPs were found in chromosome 21 in our study using single SNP GWAS, the regional heritability analysis did highlight the nearby region between 8 – 10.5 Mb as explaining over 13 % of the genetic variation of the trait. Our regional heritability analysis (RHM) also identified two additional QTL explaining a significant amount of the genetic variance, but only one of them detected in our single SNP GWAS. RHM analyses and other similar approaches use the information of several consecutive SNPs, increasing the statistical power and reliability of association mapping (Riggio and Pong-Wong, 2014; Shirali et al. 2016), which consequently should result in higher repeatability and concordance between genetic association studies. Accordingly, in our study the RHM analysis arguably located the previously detected QTL (Correa et al. 2017a), while the single SNP GWAS failed to do so.

Discovering the causal variants underlying QTL is a very challenging task, and a result very few causative variants have been identified to date. The first problem lies with the large regions that have to be investigated, since narrowing the QTL is extremely difficult due to reduced recombination and high linkage disequilibrium along large regions of the genome (e.g. Tsai et al. 2016). Further, despite the simplicity of identifying most or all variants within a region using WGS, prioritising those variants is challenging with the current status of annotation of the Atlantic salmon genome, which has 48,775 protein coding genes and 97,546 mRNAs (Lien et al. 2016). Putative non-synonymous and even premature stop mutations appear relatively frequently, probably indicating a significant proportion of pseudogenes, and therefore hindering our ability to prioritize functional mutations. Further, in complex traits, a high proportion of causative mutations are located in regulatory elements (Keane et al. 2011; Albert and Kruglyak, 2015), which are difficult to evaluate without comprehensive genome annotation using assays that can identify such regions. In this sense, the outputs of the Functional Annotation of All Salmonid Genomes (FAASG; Macqueen et al. 2017) initiative should contribute to prioritisation of intergenic SNPs through the characterization of functional regulatory elements in salmonid genomes. Complementary, differential allelic expression (DAE) and expression QTL (eQTL) can be an effective route to identify functional candidates (Gamazon et al. 2018). The caveat of DAE and eQTL is that gene expression is quite commonly restricted to specific tissues. A GTex-like project (GTEx consortium 2017) in salmonids could also facilitate the discovery of functional variants underlying QTL.

Despite these limitations, two genes were identified that are strong candidates for the QTLs in chromosome 3 and 21, TOB1 and STK17B respectively. Coho salmon, a salmonid species considered resistant to sea lice, shows pronounced epithelial hyperplasia and cellular infiltration two days after sea lice attachment, and wound healing combined with a strong Th2 immune response has been suggested as the mechanism of resistance (Braden et al. 2015). TOB1 and STK17B have been previously associated with cell proliferation and T cell regulation. TOB1 is an antiproliferative protein which is ubiquitously expressed in several species (Baranzini, 2014), and inhibits T cell proliferation in humans (Tzachanis et al. 2001). TOB1 down-regulation in response to sea lice attachment suggests that this gene plays a relevant role in the Atlantic salmon response to the parasite, and the detected putative premature stop codon mutation may be concordant with faster wound healing and T cell proliferation. STK17B, also known as DRAK2, has also been connected to T cell function (Honey 2005, Gatzka et al. 2009) and to proliferation in cancer (Yang et al. 2012; Lan et al. 2018). STK17B contains a non-synonymous mutation and marked up-regulation in response to sea lice in salmon skin. In addition to these two strong functional candidates, the allelic differential expression analysis also revealed an uncharacterized protein regulated by cis-polymorphisms in the QTL region in chromosome 3. These three genes and their mutations deserve further attention in follow-up studies aimed to increase resistance to sea lice in Atlantic salmon. Such follow up studies could include further functional annotation of QTL regions using chromatin accessibility assays to identify genomic regions potentially impacting on the binding of transcription factors or enhancers. Genome editing approaches (such as CRISPR-Cas9) could be applied to test hypotheses relating to modification of gene function or expression caused by coding or putative cis-acting regulatory variants in cell culture, or ultimately to perform targeted perturbation of the QTL regions and assess the consequences on host resistance to sea lice *in vivo*.

## CONCLUSIONS

Host resistance to sea lice in this Chilean commercial population is moderately heritable (h^2^ = 0.28) and shows a polygenic architecture, albeit with at least three QTL of moderate effect on chromosomes 3, 18 and 21 (7.8 to 13.4 % of the genetic variation). Growth during infestation also has a significant genetic component (h^2^ = 0.24), and its genetic architecture is clearly polygenic, with QTL of small effect distributed along many genomic regions. The three QTL affecting lice density were further investigated by integrating RNA-Seq and WGS data, together with a literature search. A putative premature stop codon within TOB1, an anti-proliferative protein, seems a plausible candidate to explain the QTL in chromosome 3. Alternatively, an uncharacterized protein on the same QTL region displayed differential allelic expression, and which may form a suitable target for further functional studies. STK17B, functionally connected to proliferation and T cell function, is a plausible candidate for the QTL in chromosome 21. It is evident that even when all variants in a QTL region are discovered, that shortlisting and prioritising the potential causative variants underlying QTL is challenging. However, the impending availability of more complete functional genome annotation and eQTL data is likely to assist this process, thereby helping to elucidate the functional genetic basis of complex traits in aquaculture species.

### Data Availability Statement

The RNA-Seq raw reads have been deposited in NCBI’s Sequence Read Archive (SRA)under Accession No. SRP100978, and the results have been published in Robledo et al. (2018). The WGS raw reads have been deposited in NCBI’s Sequence Read Archive (SRA) under Accession No. SRP106943. The imputed genotypes and corresponding SNP positions are available in Data Sheet 1 (compressed file, GenABEL .ped and .map files), and phenotypes of the challenged animals are available in Supplementary Table 2.

### Ethics approval and consent to participate

The lice challenge experiments were performed under local and national regulatory systems and were approved by the Animal Bioethics Committee of the Faculty of Veterinary and Animal Sciences of the University of Chile (Santiago, Chile), Certificate N° 01-2016, which based its decision on the Council for International Organizations of Medical Sciences (CIOMS) standards, in accordance with the Chilean standard NCh-324-2011.

## Authors contributions

RH, JY, and DR were responsible for the concept and design of this work and drafted the manuscript. JP was responsible for the disease challenge experiment. AB managed the collection of the samples. AG performed the molecular biology experiments. DR performed bioinformatic and statistical analyses. All authors read and approved the final manuscript.

## Acknowledgements

The authors would like to thank the contribution of Benchmark Genetics Chile and Salmones Chaicas for providing the biological material and phenotypic records of the experimental challenges. We would also like to thank Benchmark Genetics Chile for providing high-density genotypes of full-sibs of our experimental population for imputation.

*Funding statement*
This work was supported by an RCUK-CONICYT grant (BB/N024044/1) and Institute Strategic Funding Grants to The Roslin Institute (BBS/E/D/20002172, BBS/E/D/30002275 and BBS/E/D/10002070). Edinburgh Genomics is partly supported through core grants from NERC (R8/H10/56), MRC (MR/K001744/1) and BBSRC (BB/J004243/1). Diego Robledo is supported by a Newton International Fellowship of the Royal Society (NF160037). José M. Yáñez would like to acknowledge the support from Núcleo Milenio INVASAL from Iniciativa Científica Milenio (Ministerio de Economía, Fomento y Turismo, Gobierno de Chile).
*Ethics statements*
(Authors are required to state the ethical considerations of their study in the manuscript, including for cases where the study was exempt from ethical approval procedures)
*Does the study presented in the manuscript involve human or animal subjects:* No
*Data availability statement*
Generated Statement: The datasets generated for this study can be found in NCBI’s Sequence Read Archive (SRA), SRP106943.

## REFERENCES

Aaen, S.M., Helgesen, K.O., Bakke, M., Kaur, K., Horsberg, T.E. (2015). Drug resistance in sea lice: a threat to salmonid aquaculture. Trends Parasitol. 31, 72–81.

Albert, F.W., Kruglyak, L. (2015). The role of regulatory variation in complex traits and disease. Nat. Rev. Genet. 16, 197–212.

Amin, N., van Duijn, C.M., Aulchenko, Y.S. (2007). A genomic background based method for association analysis in related individuals. PLoS ONE 2, e1274.

Aulchenko, Y.S., Ripke, S., Isaacs, A., van Duijn, C.M. (2007). GenABEL: an R library for genome-wide association analysis. Bioinformatics 23, 1294–1296.

Baranzini, S.E. (2014). The role of antiproliferative gene Tob1 in the immune system. Clin. Exp. Neuroimmunol. 5, 132–136.

Bolger, A.M., Lohse, M., Usadel, B. (2004). Trimmomatic: a flexible trimmer for Illumina sequence data. Bioinformatics 30, 2114–2120.

Braden, L.M., Koop, B.F., Jones, S.R.M. (2015). Signatures of resistance to *Lepeophtheirus salmonis* include a Th2-type response at the louse-salmon interface. Dev. Comp. Immunol. 48, 178–191.

Bravo, S., Sevatdal, S., Horsberg, T.E. (2008). Sensitivity assessment of *Caligus rogercreseyi* to emamectin benzoate in Chile. Aquaculture 282, 7–12.

Canela-Xandri, O., Law A., Gray A., Woolliams, J.A., Tenesa, A., (2015). A new tool called DISSECT for analysing large genomic data sets using a big data approach. Nat. Commun. 6, 10162.

Chen, W.M., Abecasis, G.R. (2007). Family-based association tests for genomewide association scans. Am. J. Hum. Genet. 81, 913–926

Correa, K., Lhorente, J.P., López, M.E., Bassini, L., Naswa, S., Deeb, N., et al. (2015). Genome-wide association analysis reveals loci associated with resistance against *Piscirickettsia salmonis* in two Atlantic salmon (*Salmo salar* L.) chromosomes. BMC Genomics 16, 854.

Correa, K., Lhorente, J.P., Bassini, L., López, M.E., di Genova, A., Maass, A., et al. (2017a). Genome wide association study for resistance to *Caligus rogercresseyi* in Atlantic salmon (*Salmo salar* L.) using a 50K SNP genotyping array. Aquaculture 472, 61–65.

Correa, K., Bangera, R., Figueroa, R., Lhorente, J.P., Yáñez, J.M. (2017b). The use of genomic information increases the accuracy of breeding value predictions for sea louse (*Caligus rogercresseyi*) resistance in Atlantic salmon (*Salmo salar*). Gen. Sel. Evol. 49, 15.

Costello, M.J. (2009). The global economic cost of sea lice to the salmonid farming industry. J. Fish Dis. 32, 115–118.

Daetwyler, H.D., Kemper, K.E., van der Werf, J.H.J., Hayes, B.J. (2012). Components of the accuracy of genomic prediction in a multi-breed sheep population. J. Anim. Sci. 90, 3375–3384.

Dobin, A., Davis, C.A., Schlesinger, F., Drenkow, J., Zaleski, C., Jha, S., et al. (2013). STAR: ultrafast universal RNA-seq aligner. Bioinformatics 29, 15–21.

Duggal, P., Gillanders, E.M., Holmes, T.N., Bailey-Wilson, J.E. (2008). Establishing an adjusted p-value threshold to control the family-wide type 1 error in genome wide association studies. BMC Genomics 9, 516.

Fast, M.D., Johnson, S.C., Jones, S.R.M. (2007). Differential susceptibility and the responses of pink (*Oncorhynchus gorbuscha*) and chum (*O. Keta*) salmon juveniles to infection with *Lepeophtheirus salmonis*. Dis. Aquat. Organ. 75, 229–238.

Gamazon, E.R., Segrè, A.V., van de Bunt, M., Wen, X., Xi, H.S., Hormozdiari, F., et al. (2018). Using an atlas of gene regulation across 44 human tissues to inform complex disease and trait-associated variation. Nat. Genet. 50, 956–967.

Gådin JR, van’t Hooft FM, Eriksson P, Folkersen L (2015). AllelicImbalance: an R/ bioconductor package for detecting, managing, and visualizing allele expression imbalance data from RNA sequencing. BMC Bioinformatics 16, 194.

Gatzka, M., Newton, R.H., Walsh, C.M. (2009). Altered thymic selection and increased autoimmunity caused by ectopic expression of DRAK2 during T cell development. J. Immunol. 183, 285–297.

Gharbi, K., Matthews, L., Bron, J., Roberts, R., Tinch, A., et al. (2015) The control of sea lice in Atlantic salmon by selective breeding. J. R. Soc. Interface. 12, 0574.

Gilmour, A.R., Gogel, B.J., Cullis, B.R., Thompson, R. (2014). ASReml user guide. 4th ed. Hemel Hempstead: VSN International Ltd.

Gjerde, B., Ødegård, J., Thorland, I. (2011). Estimates of genetic variation in the susceptibility of Atlantic salmon (*Salmo salar*) to the salmon louse *Lepeophtheirus salmonis*. Aquaculture 314, 66–72.

GTEx Consortium (2017). Genetic effect on gene expression across human tissues. Nature 550, 204–210.

Honey, K. (2005). T-cell signalling: DRAK2 puts the brakes on T-cell responses. Nat. Rev. Immunol. 5, 98.

Houston, R.D. (2017). Future directions in breeding for disease resistance in aquaculture species. R. Bras. Zootec. 46, 545–551.

Jackson, D., Moberg, O., Stenevik Djupevåg, E.M., Kane, F., Hareide, H. (2017). The drivers of sea lice management policies and how best to integrate them into a risk management strategy: an ecosystem approach to sea lice management. J. Fish. Dis. 41, 927–933.

Jónsdóttir, H., Bron, J.E., Wootten, R., Turnbull, J.F. (1992). The histopathology associated with the pre-adult and adult stages of *Lepeophtheirus salmonis* on the Atlantic salmon, *Salmo salar* L. J Fish Dis. 15, 521–527.

Keane, T.M., Goodstadt, L., Danecek, P., White, M.A., Wong, K., Yalcin, B., et al. (2011). Mouse genomic variation and its effect on phenotypes and gene regulation. Nature 477, 289–294.

Kofler, R., Pandey, R.V., Schlötterer, C. (2011). PoPoolation2: identifying differentiation between populations using sequencing of pooled DNA samples (Pool-Seq). Bioinformatics 27, 3435–3436.

Kolstad, K., Heuch, P.A., Gjerde, B., Gjedrem, T., Salte, R. (2005) Genetic variation in resistance of Atlantic salmon (*Salmo salar*) to the salmon louse *Lepeophtheirus salmonis*. Aquaculture 247, 145–51.

Lan, Y., Han, J., Wang, Y., Wang, J., Yang, G., Li, K., et al. (2018). STK17B promotes carcinogenesis and metastasis via AKT / GSK-3β / Snail signalling in hepatocellular carcinoma. Cell Death Dis. 9, 236.

Lhorente, J.P., Gallardo, J.A., Villanueva, B., Araya, A.M., Torrealba, D.A., Toledo, X.E., et al. (2012). Quantitative genetic basis for resistance to *Caligus rogercresseyi* sea lice in a breeding population of Atlantic salmon (*Salmo salar*). Aquaculture 324-325, 55–59.

Lhorente, J.P., Gallardo, J.A., Villanueva, B., Carabaño, M.J., Neira, R. (2014). Disease resistance in Atlantic salmon (*Salmo salar*): coinfection of the intracellular bacterial pathogen *Piscirickettsia salmonis* and the sea louse *Caligus rogercresseyi*. PLoS One 9, e95397.

Li, H., Handsaker, B., Wysoker, A., Fennell, T., Ruan, J., Homer, N., et al. (2009). The sequence alignment/map (SAM) format and SAMtools. Bioinformatics 25, 2078–2079.

Li, H. (2013). Aligning sequence reads, clone sequences and assembly contigs with BWA-MEM. arXiv:1303.3997v1 [q-bio.GN].

Lien, S., Koop, B.F., Sandve, S.R., Miller, J.R., Kent, M.P., Nome, T., et al. (2016). The Atlantic salmon genome provides insights into rediploidization. Nature 533, 200–205.

Macqueen, D.J., Primmer, C.R., Houston, R.D., Nowak, B.F., Bernatchez, L., Bergseth, S., et al. (2017). Functional annotation of all salmonid genomes (FAASG): an international initiative supporting future salmonid research, conservation and aquaculture. BMC Genomics 18, 484.

Matsuda, S., Kawamura-Tsuzuku, J., Ohsugi, M., Yoshida, M., Emi, M., Nakamura, Y., et al. (1996). Tob, a novel protein that interacts with p185erbB2, is associated with anti-proliferative activity. Oncogene 12: 705–713.

Meuwissen, T.H., Hayes, B.J., Goddard, M.E. (2001). Prediction of total genetic value using genome-wide dense marker maps. Genetics 157, 1819–1829.

Nagamine, Y., Pong-Wong, R., Navarro, P., Vitart, V., Hayward, C., Rudan, I., et al., 2012. Localising loci underlying complex trait variation using regional genomic relationship mapping. PLoS ONE 7: e46501

Ødegård, J., Moen, T., Santi, N., Korsvoll, S.A., Kjøglum, S., Meuwissen, T.H.E. (2014). Genomic prediction in an admixed population of Atlantic salmon (*Salmo salar*). Frontiers in Genetics 5, 402.

Pulgar, R., Hödar, C., Travisany, D., Zuñiga, A., Domínguez, C., Maass, A., et al. (2015). Transcriptional response of Atlantic salmon families to *Piscirickettsia salmonis* infection highlights the relevance of the iron-deprivation defence system. BMC Genomics 16, 495.

Riggio, V., and Pong-Wong, R. (2014). Regional Heritability Mapping to identify loci underlying genetic variation of complex traits. BMC Proc. 8, S3.

Sargolzaei, M., Chesnais, J.P., Schenkel, F.S. (2014). A new approach for efficient genotype imputation using information from relatives. BMC Genomics 15, 478.

Shirali, M., Pong-Wong, R., Navarro, P., Knott, S., Hayward, C., et al. (2016) Regional heritability mapping method helps explain missing heritability of blood lipid traits in isolated populations. Heredity 116, 333–338.

Sutherland, B.J.G., Koczka, K.W., Yasuike, M., Jantzen, S.G., Yazawa, R., Koop, B.F., et al. (2014). Comparative transcriptomics of Atlantic *Salmo salar*, chum *Oncorhynchus keta* and pink salmon *O. gorbuscha* during infections with salmon lice *Lepeophtheirus salmonis*. BMC Genomics 15, 200.

Robledo, D., Matika, O., Hamilton, A., Houston, R.D. (2018). Genome-wide association and genomic selection for resistance to amoebic gill disease in Atlantic salmon. G3 8, 1195–1203.

Robledo, D., Gutiérrez, A.P., Barría, A., Yáñez, J.M., Houston, R.D. (2018). Gene expression response to sea lice in Atlantic salmon skin: RNA sequencing comparison between resistant and susceptible animals. Front. Genet. 9, 287.

Taylor, R.S., Kube, P.D., Muller, W.J., Elliott, N.G. (2009). Genetic variation of gross gill pathology and survival of Atlantic salmon (*Salmo salar* L.) during natural amoebic gill disease challenge. Aquaculture 294, 172–179.

Tsai, H.Y., Hamilton, A., Tinch, A.E., Guy, D.R., Bron, J.E., Taggart, J.B., et al. (2016) Genomic prediction of host resistance to sea lice in farmed Atlantic salmon populations. Gen. Sel. Evol. 48, 47.

Tzachanis, D., Freeman, G.J., Hirano, N., van Puijenbroek, A.A.F.L., Delfs, M.W., Berezovskaya, A., et al. (2001) Tob is a negative regulator of activation that is expressed in anergic and quiescent T cells. Nat. Immunol., 2, 1174–1182.

Uemoto, Y., Pong-Wong, R., Navarro, P., Vitart, V., Hayward, C., et al. (2013). The power of regional heritability analysis for rare and common variant detection: simulations and application to eye biometrical traits. Front. Genet. 4, 232.

Wang, Y., Lin, G., Li, C., Stothard, P. (2016). Genotype imputation methods and their effects on genomic predictions in cattle. Springer Sci. Rev. 4, 79.

Yáñez, J.M., Lhorente, J.P., Bassini, L.N., Oyarzún, M., Neira, R., Newman, S. (2014). Genetic co-variation between resistance against both *Caligus rogercresseyi* and *Piscirickettsia salmonis*, and body weight in Atlantic salmon (*Salmo salar*). Aquaculture 433, 295–298.

Yáñez, J.M., Naswa, S., López, M.E., Bassini, L., Correa, K., Gilbey, J., et al. (2016). Genomewide single nucleotide polymorphism discovery in Atlantic salmon (*Salmo salar*): validation in wild and farmed American and European populations. Mol. Ecol. Res. 16, 1002–1011.

Yang, K.M., Kim, W., Gim, J., Weist, B.M., Jung, Y., Hyun, J.S., et al. (2012). DRAK2 participates in a negative feedback loop to control TGF-β/Smads signalling by binding to type I TGF-β receptor. Cell Rep. 2, 1286–1299.

